# Feedback to deep layers in human V1 during perceptual filling-in

**DOI:** 10.64898/2026.04.17.719145

**Authors:** Kenshu Koiso, Anna Razafindrahaba, Vincent Van De Ven, Mark J. Roberts, Federico De Martino, Peter De Weerd

## Abstract

Visual surface perception is a fundamental aspect of vision, yet its neural implementation remains poorly understood. Troxler’s perceptual filling-in paradigm provides a tractable illusion for studying surface perception, in which a peripheral figure becomes perceptually assimilated into the surrounding background after a period of sustained fixation. Although neural correlates of this phenomenon have been reported in early visual cortex, the underlying mechanisms, particularly the contribution of feedback signaling, remain unresolved. Here we use ultra-high-field (7T) layer-fMRI to investigate perceptual filling-in in the human visual cortex. While experimentally controlling perceptual filling-in, we measured GE-BOLD responses in ten participants. Analyses across cortical depth in the independently localized figure representation in primary visual cortex (V1) revealed neural correlates of filling-in in deep cortical layers, which are associated with feedback input. These findings provide evidence that perceptual filling-in and visual surface perception in general are supported by feedback signals to early visual cortex.

## Introduction

In natural visual scenes, surface boundaries are most likely to correspond to strong luminance and other contrasts, whereas surface interiors are ordinarily characterized by weak contrasts and homogenous stimulation (Lowet et al., 2015; Marr, 1982; Simoncelli & Olshausen, 2001). The center-surround organization of retinal ganglion cell receptive fields (Kuffler, 1953) renders the early visual system highly sensitive to local contrast at surface boundaries, but comparatively insensitive to uniform surface regions. This organization implies that the perception of extended surface properties cannot be fully explained by feedforward processing alone, and instead likely depends on cortical mechanisms that reconstruct, or fill-in, surface representations beyond the information directly available in the retinal input (De Weerd et al., 2006; Pessoa et al., 1998; Walls, 1954; Weil & Rees, 2011).

To isolate cortical contributions to surface filling-in, visual illusions have been used that elicit surface percepts in the absence of corresponding physical evidence (Komatsu, 2006; Spillmann & Dresp, 1995). The overall body of evidence from different experimental paradigms supports contributions from both striate and extrastriate cortex to surface perception. V1 neural responses have been shown to correlate positively with brightness induction in cats (Rossi et al., 1996). Other neurophysiological studies using the Craik O’Brain Cornsweet illusion (Cornsweet, 1970) have shown activity increases associated with enhanced surface brightness in cat area 18 (Hung et al., 2001, 2007) and in monkey V2 (Roe et al., 2005). Komatsu et al., (1996, 2002) observed neural responses during surface filling-in across the blind spot, and De Weerd et al., (1995) recorded response increases in monkey V2 and V3 associated with texture filling-in in a Troxler paradigm. In humans, functional magnetic resonance imaging (fMRI) studies using Kanizsa figures (Kanizsa, 1976; Kok et al., 2016; Kok & de Lange, 2014) and neon color spreading (Sasaki & Watanabe, 2004; Van Tuijl, 1975) have reported activity related to illusory surface perception. Brightness induction in other human fMRI studies revealed neural correlates in V1 (Malik & Boyaci, 2024; Pereverzeva & Murray, 2008) in V2 (van de Ven et al., 2012), or in V1, V2 and V3 (Salmela & Vanni, 2013). fMRI studies using the Troxler paradigm, have reported diverse findings. Hsieh and Tse (2010) demonstrated a multivariate representation in V1 of perceived color mixing (Hsieh & Tse, 2010), whereas Mendola et al. (2006) reported decreased V1/V2 activity and increased V3/V4 activity during fading of a disk of slightly higher luminance than its background (Mendola et al., 2006).

Advancing our understanding of surface-related activity in retinotopic areas requires addressing fundamental questions about its underlying mechanisms. Since retinotopic surface representations are unlikely to arise from feedforward input alone, feedback comes to the fore as a plausible source of surface-related activation in early visual areas (Roelfsema et al., 2002). Anatomical (Felleman & Van Essen, 1991; Markov et al., 2014; Vezoli et al., 2021) and neurophysiological studies (Roberts et al., 2013; Self et al., 2013; Van Kerkoerle et al., 2017) have established that feedforward projections predominately target layer IV in higher-level areas, whereas feedback projections target both deep and superficial layers in lower-level cortex, while avoiding the middle layer. Moreover, short-range feedback preferentially targets the superficial layers and long-range feedback the deep layers (Markov & Kennedy, 2013; Vezoli et al., 2021). Therefore, if activity in a retinotopic cortical region related to surface perception were predominant in deep or superficial layers, this would suggest a feedback contribution originating from respectively remote or more nearby sources. There has so far been only a single ultra-high-field fMRI study in which inducing the percept of an illusory surface was associated with feedback targeting deep layers in V1 (Kok et al., 2016). The illusory surface percept in their study was induced by a Kanizsa figure. To assess whether Kok et al.’s (2016) finding revealed a fundamental aspect of surface perception, or a finding limited to Kanizsa figures, we tested whether deep-layer feedback contributions generalize to surface filling-in generated with experimental stimuli other than Kanizsa figures.

To accomplish this goal, we used the Troxler filling-in paradigm (Troxler, 1804), which provides a powerful illusory percept of surface filling-in. In our version of this paradigm, which closely matches that used in earlier monkey neurophysiological work (De Weerd et al., 1995), we surrounded a grey square with a dynamic line texture (De Weerd et al., 1995, 1998), and instructed participants to fixate a point away from the figure. Maintained fixation induced the physically grey figure to become perceptually filled-in by the surrounding texture background (De Weerd et al., 1998; Ramachandran & Gregory, 1991; Spillmann & Kurtenbach, 1992). Because perceptual filling-in occurs spontaneously in the absence of physical changes to the stimulus, this paradigm is well-suited for isolating feedback-related neural processes from feedforward influences related to stimulus input. Here, we used ultra-high-field (7T) laminar-specific fMRI in human participants to investigate the laminar mechanisms in early visual cortex underlying texture-based perceptual filling-in. We hypothesized that perceptual filling-in would be associated with activity changes in deep and /or superficial layers of early visual cortex consistent with a feedback-driven mechanism.

## Results

Dynamic textures of white vertical bars on a black background were presented in 30 s blocks. Throughout each block, participants fixated a point in the bottom-right corner while a 2.75° × 2.75° grey square, equiluminant with the surrounding texture, was displayed at 5.5° eccentricity (Figure 1). In the filling-in enabling condition (FIE; Figure 1A), the dynamic texture was presented continuously. In the filling-in prevention condition (FIP; Figure 1B), the dynamic texture background was replaced regularly (once per second) by 300 ms of grey frames, to counteract filling-in. We additionally included a full-field texture (TEX) condition (Figure 1C) to test the possibility that activity observed during perceptual filling-in in the FIE condition would approach the activity level driven by physical texture. Blocks of FIE, FIP, and TEX alternated in the fMRI experiment with rest periods, in which a uniform grey screen was presented while fixation was maintained (Figure 1D). During both FIE and FIP blocks, participants used keyboard button presses to rate the perceived degree of filling-in on a four-point scale, from minimal filling-in near the figure borders (level 1) to complete filling-in (level 4; Figure 1E).

**Figure 1.**
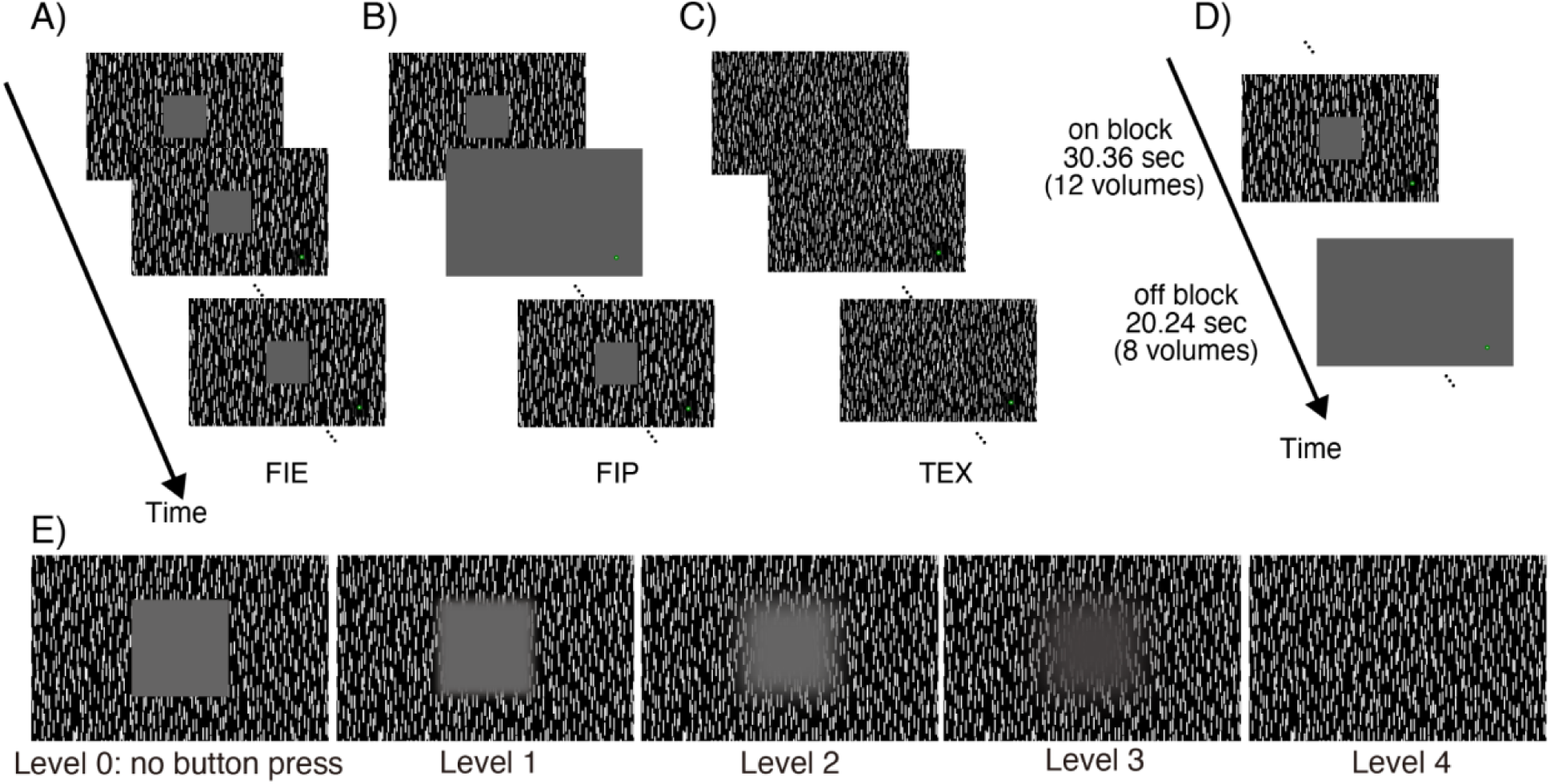
Experimental design. **(A-C)** Each fMRI run included three conditions: the filling-in enabling condition (FIE), the filling-in prevention condition (FIP), and the full-field texture condition (TEX). In FIE, a 2.75° x 2.75° equiluminant grey square was embedded in a dynamic texture background, while participants fixated a spot at the right-hand bottom of the screen 5.5° away from the center of the square. FIP used the same configuration but intermittently replaced the texture with isoluminant grey for 0.3 s at a frequency of 1 Hz. In TEX, only the dynamic texture and the fixation spot were shown. The white fixation spot had a diameter of 0.1°, which was placed in the middle of a green disk of 0.3° diameter. The fixation spot and its green surround were centered in the middle of a circular region of 1.5° diameter devoid of dynamic line elements. Mean luminance was 5 cd/m^2^ throughout the display in all stimulus conditions. **(D)** The fMRI session interleaved 30.36 s stimulation blocks with 20.24 s rest periods. FIE blocks and FIP blocks appeared in a balanced order (four per condition per run). TEX blocks were presented in separate runs. **(E)** Participants reported perceived filling-in according to a 5-level scale using four buttons. Level 0 corresponded to the absence of filling-in (no button press). Further levels ranged from 1 (blurred edges) and 2-3 (texture perceptually flowing into the figure) to level 4 (complete filling-in). These levels were assigned to button presses from index (level 1) to little finger (level 4), and a specific button press was maintained as long as the corresponding percept lasted. To provide a criterion for judging the different levels of filling-in, participants were shown the five digital images associated to levels 0 to 4 before the start of the experiment. Upon questioning after the experiment, the participants confirmed that the scheme with four levels of filling-in captured their subjective experience.

Our data showed robust behavioral evidence for perceptual filling-in. Figure 2A shows behavioral reports obtained during fMRI scanning from three representative participants (see Supplementary Figure S1 for all participants), comparing data from the FIE (left) and the FIP (right) conditions. After several seconds of stimulus presentation, participants showed considerable filling-in in the FIE condition (darker grey tones represent stronger filling-in) and comparatively limited filling-in in the FIP condition. This difference was representative for the group of participants (Figure 2B), in which the average level of filling-in plateaued towards the end of the FIE condition, reaching an average level of 2.5, as opposed to an average level of 0.5 in the FIP condition. Figures 2B and 2C show behavioral data from 9 of the 10 scanned participants, because in one of them behavioral data was lacking due to a technical failure (for details, see Supplementary Figure S1). A paired Wilcoxon signed-rank test showed a significant difference between FIE and FIP conditions starting 7.5 s after stimulus onset (Figure 2B, stars show significant differences per time point at p < 0.05, Bonferroni corrected). Within the range of times in which there was significantly more filling-in in FIE compared to FIP conditions (indicated by stars in Figure 2B), late time points (22.77 – 30.36 s) consistently showed more filling-in than early time points (7.59 – 17.71 s) (Figure 2C, two-tailed paired t-test, t(8) = 4.87, p = 0.001).

**Figure 2.**
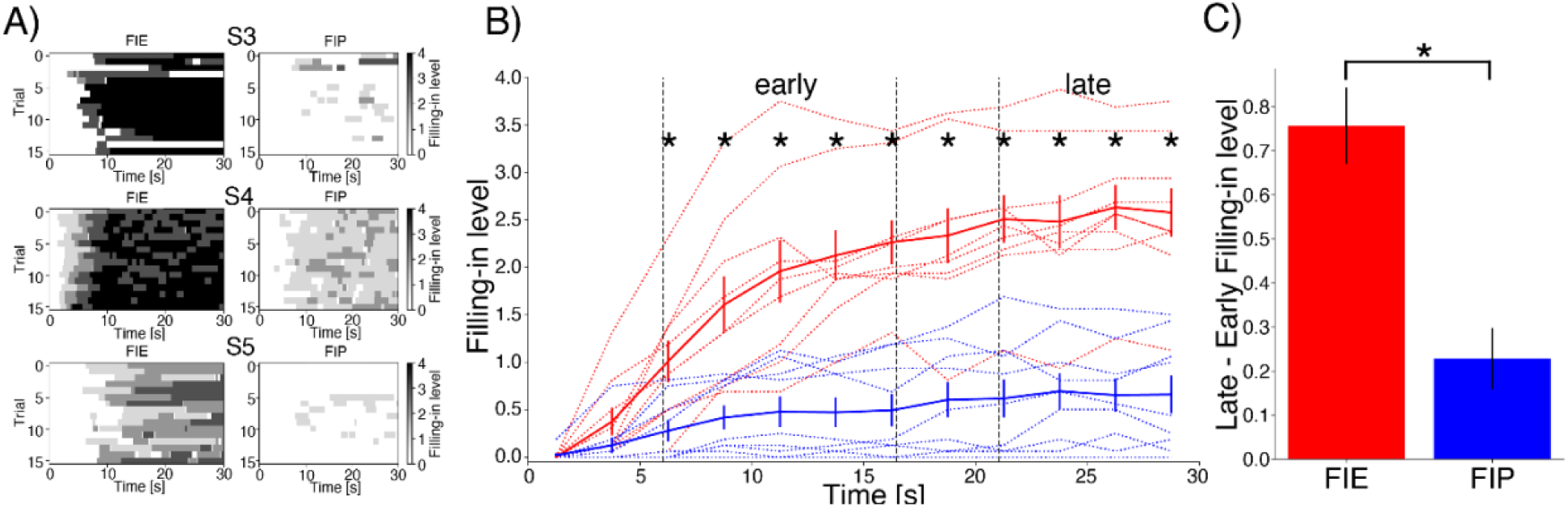
Behavioral filling-in reports. **(A)** Examples of individual filling-in reports in three participants. Within each panel, rows represent individual trials, while the color (light to dark grey) represents the level of the reported filling-in percept. **(B)** Filling-in level time course. Average across participants is shown in solid lines and individual participant data is shown in thin dotted lines. Stars show statistical significance between conditions (paired Wilcoxon signed-rank test, p < 0.05, Bonferroni corrected) **(C)** Increase in filling-in level from early to late periods (level in early period is subtracted from level in late period) in FIE and FIP conditions. Stars show statistical significance between conditions (paired t-test, t(8) = 4.87, p = 0.001). Error bars represent standard errors of the mean across participants in B and C.

The strong psychophysical difference between FIE and FIP conditions afforded us with the degree of experimental control required to isolate a signal related to perceptual filling-in by contrasting fMRI activity in the FIE and FIP conditions. In our experiments, we chose the largest figure that still allowed robust filling-in (2.75° x 2.75° at an eccentricity of 5.5°, based on pilot behavioral tests). However, because the V1 population receptive field (pRF) size at the figure’s eccentricity (about 1°; Dumoulin & Wandell, 2008) is relatively large compared to the chosen figure size, we anticipated that feedforward input from the background would contribute partially to activity measured in the figure’s ROI in V1 as well as in extrastriate cortex. We furthermore anticipated that any feedforward influences would be stable through the trial, whereas feedback influences related to the percept of filling-in would increase through the trial, possibly approaching the level of activity elicited by the physical texture. Therefore, our analysis of fMRI signal focused on changes in signal strength from the early to the late period of the trial.

The visual cortical response to perceptual filling-in and the contribution of feedback were tested in the region of low-level cortex that represented the area of the figure. To define a retinotopic region of interest (ROI) corresponding to the figure’s representation, we collected independent localizer runs. From the localizer runs, we were only able to reliably determine a ROI in area V1 in all participants. In 6 of the 10 participants, we were also able to identify small ROIs in area V2, however given their small size we did not pursue further analysis in these regions (see Methods and Supplementary Figure S2 for more details). Within the individually defined V1 ROIs, we extracted event-related averages (normalized to percent signal change) for each participant and condition of interest. We then compared fMRI signal in the early time window, showing limited perceptual filling-in (7.59 – 17.71 s), with a late time window showing strong filling-in (22.77-32.89 s).

We had expected an increase in activity within the V1 ROI specifically in the FIE but not the FIP condition, by analogy with the perceptual data shown in Figure 2. However, the fMRI signal combined over cortical depths showed a small, but significant signal increase in both conditions (see Supplementary Figure S3, and Discussion). An ANOVA revealed a main effect of time, without a time x condition interaction (time, F(1,36) = 21.47, p < 0.001; condition, F(1,36) = 0.54, p = 0.46; interaction, F(1, 36) = 0.12, p = 0.72). Notably, in the present and in further analyses, we excluded the TEX condition. This condition had been included because we had expected that the filling-in-related increase would approach a level similar to that driven by physical texture inside the ROI. The magnitude of the filling-in-related signal, however, was very small compared to the level of activity produced by physical texture (Supplementary Figure S3). This rendered the planned comparisons of the activity level during perceptual filling-in (FIE) with the activity level elicited during physical filling-in (TEX) uninteresting, which led to the removal of the TEX condition from analysis.

The depth-averaged fMRI signal increase in both FIE and FIP conditions might have obscured meaningful signal differences among these conditions at the level of cortical layers. We reasoned that during the FIE condition, which generated the percept of filling-in, signal increases might have been dominated by increased feedback and thus would be more pronounced in superficial and deep compared to middle layers. The FIP condition, on the other hand, might have been associated predominantly with feedforward drive, and thus any fMRI signal increase due to feedforward influences would be more pronounced in middle compared to superficial and deep layers. Hence, we hypothesized a relatively larger contribution of deep and/or superficial layers to the average signal increase in the FIE condition, as opposed to a relatively larger contribution of the middle layer to the signal increase in the FIP condition.

To test this hypothesis, we segmented the cortex into superficial, middle and deep layer compartments (see Methods) and calculated fMRI signal increases from early to late time periods of the event-related average response in the middle and feedback (deep and superficial) layers separately. We obtained a *feedback index* by contrasting activity in feedback (deep and superficial) and middle layers (see Methods). A positive index points to greater increases in feedback layers compared to the middle layer, whereas a negative index points to a greater increase in the middle layer compared to the feedback layers. In agreement with our hypothesis, we found a positive feedback index in the FIE condition, a negative index in the FIP condition, and a significant difference between the two conditions (paired t-test, t(9) = 3.73, p = 0.005). This indicates that the V1 signal related to perceptual filling was driven by feedback influences (Figure 3A).

**Figure 3.**
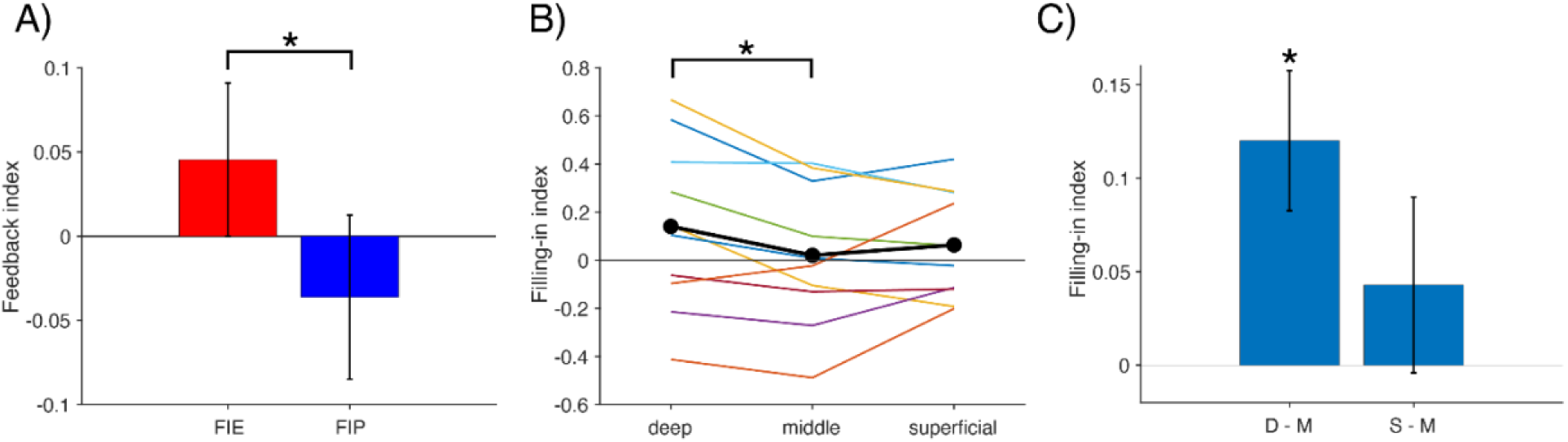
Laminar analysis. **(A)** Quantification of the feedback index (percent signal increase in deep and superficial layers minus percent signal increase in middle layer) for FIE and FIP conditions (mean value across participants, error bars show standard error). The feedback index reflects differences among layers in the magnitude of activity increases over time (see Supplementary Figure S3), with stronger increases in feedback layers in FIE and stronger increase in the middle layer in the FIP condition. **(B)** Difference in percent signal increase (late minus early time window) between FIE and FIP conditions (FIP minus FIE) across cortical depths (deep, middle and superficial) in all individual participants. In the resulting ‘Filling-in Index’, a positive value means a larger activity increase during the FIE condition than in the FIP condition. Colored lines indicate individual participants. **(C)** Same as in B for deep and superficial layers after contrasting to the middle layer: The filling-in index was normalized by subtracting the index in the middle layer from that in deep (D - M) and superficial layers (S - M) in each participant.

We then carried out a complementary analysis to compare the magnitude of signal increases in FIE and FIP conditions across cortical depths using a *filling-in index*. The filling-in index quantifies how much more activity increase there was in the FIE compared to FIP condition. Thus, we expected a positive filling-in index in the superficial and/or deep feedback layers, indicating a larger activity increase in FIE than in FIP condition. In the middle layer, we expected a comparatively smaller filling-in index, which might have reached negative values if signal increases in the middle layer were more pronounced in the FIP than in the FIE condition. The data revealed large inter-individual differences in the magnitude of the filling-in index (Figure 3B). Nevertheless, we found that overall the filling-in index was significantly different between deep and middle layers (Figure 3B and 3C; paired t-test, t(9) = 3.21, p = 0.011), whereas this index was not significantly different between superficial and middle layers (paired t-test, t(9) = 0.91, p = 0.39). A larger filling-in index in deep compared to middle layers but not in superficial compared to middle layers indicates that feedback contributions during perceptual filling-in may target deep layers more prominently.

The results in Figure 3A-C were computed using all cortical locations activated by the figure presentation during the localizer mapping. During this mapping, the figure was covered in 8 phase-encoding steps (details in Methods) from its center (phase-encoding step 1) to its edge (phase-encoding step 8). We repeated the previous analyses considering progressively expanding subsets of cortical regions: voxel groupings including steps 1 to 4, steps 1 to 5, steps 1 to 6, steps 1 to 7, or steps 1 to 8 were investigated. The difference in the feedback index remained significant for vertex groupings concatenating steps 1 to 6, steps 1 to 7, and steps 1 to 8, but not for the voxel groupings limited to the ROI center (1 to 4 and 1 to 5). This argues against the idea that the magnitude of the feedback index was limited by, or strongly determined by, our choice to include all ROI vertices (steps 1 to 8) into the computation of this index (Supplementary Figure S4A). Furthermore, the finding that expanding the 1-6 step vertex grouping to include steps 7 and 8 did not strongly alter the pattern of results suggests that the signal we captured was dominated by a correlate of perceptual filling-in rather than by an adapting response to the figure-ground boundary. For the filling-index, we evaluated the significance for the same voxel groupings as for the feedback index and found that beyond the significant effect in the standard group (phase encoding steps 1-8), the index remained significant also for the 1-7 steps vertex group, but not for the other groups (Supplementary Figure S4B). We also tested how robust our results were to different potential choices for the early and late time windows. We used a briefer early (7.59 – 15.18 s) and late time window (25.30 - 32.89 s) and moved the early window closer to the late window in several steps (1 TR shift per step) while the late window’s position remained fixed. We found both feedback index and filling-in index were robust against variation in the choice of the early window (see Supplementary Figure S5). In conclusion, the central finding of our work is a small but robust signal increase in deep V1 layers related to perceptual filling-in.

## Discussion

In this study, we measured the neural correlates of perceptual filling-in in V1. Using 7T fMRI, we compared responses from the representation of a physically grey square under conditions that either enabled perceptual filling-in of the figure by surrounding dynamic texture (FIE condition) or prevented filling-in (FIP condition). We hypothesized activity in the retinotopic representation of the figure to increase in the FIE condition, as opposed to activity in the FIP condition. Contrary to our expectations, we did not observe an increase in V1 activity limited to the FIE condition, instead, both the FIE and FIP conditions showed a small activity increase over time. However, when comparing activity among V1 layers, we consistently found a steeper activity increase in the FIE than in the FIP condition in the deep layers. This layer-specific effect is consistent with the interpretation that perceptual filling-in engages feedback signals targeting deep layers of V1.

De Weerd et al. (1998) suggested that high-level cortical areas may play a role in surface perception. In line with that suggestion, neurophysiological (Roelfsema et al., 2002; Self et al., 2013; Van Kerkoerle et al., 2014) and optogenetics studies (Kirchberger et al., 2023; Pak et al., 2020) have confirmed the critical contributions of higher-level mechanisms to figure-ground segregation. Moreover, Wokke et al. (2013) demonstrated a critical role of lateral occipital cortex (LOC) in the perception of the Kanizsa illusion using transcranial magnetic stimulation (TMS). In the Troxler paradigm, continued stabilization of the stimulus and ensuing failure of figure-ground segregation may lead to a change in feedback signal from high-level areas to the retinotopic representations of the figure in low-level areas, which may underlie the percept of filling-in.

Feedback connections enter both superficial and deep layers, whereby short-range feedback preferentially targets the superficial layers and long-range feedback the deep layers (Markov & Kennedy, 2013; Vezoli et al., 2021). In a human 7T fMRI study using a Kanizsa figure (Kok et al., 2016), a monkey neurophysiological study of the blind-spot (Komatsu et al., 2002), as well as in the present study, the perceived changes in brightness or texture during surface filling-in induced signal in deep V1 layers, consistent with long-range feedback originating in high-level visual areas. By contrast, other 7T fMRI studies have reported activity in superficial layers of the V1 representation of a grey surface carrying contextual scene-related information (Muckli et al., 2015) or motion-related information (Marquardt et al., 2020). These paradigms without visible changes in the surface’s appearance therefore seem associated with short-range feedback to V1. However, although less numerous, superficial layers in V1 also receive long-range feedback connections (Rockland et al., 1994), and deep layers in V1 also receive short-range feedback connections (Anderson & Martin, 2009). The latter, in combination with the correlate of perceptual filling-in reported in V2 and V3 in a neurophysiological study (De Weerd et al., 1995), suggests that short-range feedback from extrastriate to striate cortex may also have contributed to the filling-in related fMRI signal we have observed in deep V1 layers.

Although our data support a role of feedback during perceptual filling-in, we cannot establish in the current study what the specific contribution of feedback is during the emergence of this percept. Many computational models of surface perception include a spreading or diffusion process initiated and contained by boundaries as an important component in surface representation (Grossberg, 1987a, 1987b; Neumann et al., 2001; Peters et al., 2010; Shipley & Kellman, 1994). Lateral spreading of neural activity carrying information about surface features could be accomplished via lateral connections within early areas (Lund et al., 2003; McManus et al., 2008). In the Troxler paradigm, retinal stabilization of the figure is thought to lead to the adaptation of boundary representations, followed by an invasion of background-related activity into the representation formerly occupied by the figure (Gerrits & Vendrik, 1970; Spillmann & De Weerd, 2003). In this context, the activity in deep layers we have observed might represent a signal from high-level regions extinguishing the weakening boundary representations, thus permitting the lateral spread of activity related to the background feature. Alternatively, the feedback signal might also directly implement the lateral spreading of information (Poort et al., 2016; Roelfsema et al., 2002). In our data, we did not have the spatial resolution to separately capture declining activity at the figure boundary that is expected to occur concurrently with activity increases related to perceptual filling-in.

The pattern of fMRI data observed here appears at odds with the neuronal spiking data recorded from monkey V1, V2 and V3 neurons with receptive fields contained within a figure surrounded by dynamic texture (De Weerd et al., 1995). This study reported that while the monkeys continually fixated a spot away from the figure, extrastriate but not V1 neurons showed increases in neural activity coinciding with the onset of perceptual filling-in in humans tested with the same stimuli. Beyond the fact that fMRI may be more sensitive in detecting subtle feedback influences than neuronal spiking recordings (Logothetis et al., 2001), there are at least three reasons that might underlie the different V1 results in their neurophysiological study and our fMRI study. First, De Weerd et al. (1995) reported data from only a small sample of V1 neurons whose layer distribution was unverified. It is therefore possible that spiking increases related to perceptual filling-in were missed as the recorded neurons might have been outside the deep layers. Second, assuming De Weerd et al. (1995) missed the V1 neurons contributing to filling-in by recording mostly cells in superficial layers, we expected a substantial fMRI signal increase in V1 associated with filling-in, mimicking their extrastriate results. Instead, we observed only a small signal increase in V1 related to perceptual filling-in. It is possible that the modest fMRI signal increase in comparison to that observed in neurophysiological recordings is due to the different conditions under which the neurophysiological and fMRI data were collected. The monkeys in De Weerd et al. (1995) did a passive fixation task while (likely) ignoring the figure, whereas the human participants in the present fMRI study reported each episode of perceptual filling-in with button presses. Attention has been reported to increase or decrease lateral interactions (Freeman et al., 2001; Herrero et al., 2008; Ito & Gilbert, 1999; Roberts & Thiele, 2008) depending on stimulus condition and eccentricity. It is therefore a possibility that in our study, perceptual filling-in (and its neural correlate) might have been counteracted by paying attention to the figure due to the instruction to report it. Further, because of their extensive training, the monkeys in De Weerd et al. (1995) likely fixated more accurately than our human participants during fMRI scanning, and accurate fixation promotes filling-in (Martinez-Conde et al., 2006). Hence, more accurate fixation and lack of attention to the figure might have led to a fast invasion of the figure by the background representation in the monkey neurophysiological study (De Weerd et al., 1995), as indexed by the strong responses in V2 and V3. Conversely, responses during perceptual filling-in in the human participants might have been much smaller, due to attention to the figure and less accurate fixation. Third, in line with the preceding argument, it is interesting to note that the average level of filling-in on the 1-4 response scale we used was 2.5. This suggests that the fMRI feedback signature related to filling-in that we detected was more related to the perceptual filling-in of the brightness level of the background than to the texture filling-in *per se* for a majority of trials and participants (see Figure 1E and Supplementary Figure S1A). This is in line with psychophysical studies showing that different surface features can perceptually fill-in with different time courses (Ramachandran & Gregory, 1991; Welchman & Harris, 2001). Further, although responses to brightness changes have been reported to range from small (Kok & de Lange, 2014; Roe et al., 2005; Sasaki & Watanabe, 2004) to considerable (Malik & Boyaci, 2024; Rossi et al., 1996), responses to a perceived switch from grey to dynamic texture could reasonably be expected to produce a more pronounced neural response increase (De Weerd et al., 1995) than what we observed. If this is correct, and the percept of filling-in in our data was driven more by a subtle change in brightness in the figure region, and less by the subjective emergence of the dynamic line texture, then this could also have contributed to our finding of a correlate of perceptual filling-in less vigorous than expected. An additional factor that may have contributed to a smaller-than-expected fMRI signal related to filling-in is related to the constraints on our stimulus imposed by the small field-of-view in the 7T MRI scanner. To ensure a sufficient frame of texture surrounding the figure, we could not place the figure very peripherally. Instead, we had to use a fairly small figure that we then placed close enough to the fixation spot to have a sizeable representation while still allowing perceptual filling-in (De Weerd et al., 1998). This unavoidable compromise had the consequence that the voxels in the figure ROI were driven to a significant extent by the texture background, rendering the detection of signal related to perceptual filling-in more difficult. The small figure ROI also made it difficult to localize the V2/V3 ROI, which prompted us to limit the present report to V1.

GE-BOLD signal, as used in the present study, is sensitive to large veins, and therefore biases signal strength towards superficial layers. Although specific models have been proposed to mitigate venous effects in the BOLD signal (Corbitt et al., 2018; Faes et al., 2025; Havlicek & Uludağ, 2020; Heinzle et al., 2016; Markuerkiaga et al., 2016; Marquardt et al., 2020; Puckett et al., 2016; Uludağ, 2023), the various parameters in these models are not easily optimized. Hence, in the present study, like others (Bergmann et al., 2024; Kok et al., 2016; Lawrence et al., 2018) we did not apply a corrective model but instead contrasted conditions of interest within layers. We suggest that this experimental approach largely addresses the concern that our conclusions may be driven by venous bias favoring signal in superficial layers. The fact that the signal enhancement of interest in our study was predominant in deep layers further supports our conclusion of a genuine feedback signature related to perceptual filling-in.

We observed a very small but statistically significant trend for fMRI signal averaged over cortical depths to increase over time, irrespective of the stimulus condition. This small signal increase in the cortex in all experimental conditions could be due to arousal slowly building up towards the end of fMRI stimulation blocks through the effects of noradrenaline and other neuromodulators (Huang et al., 2023; Reteig et al., 2019). The stronger increase of activity during perceptual filling-in the FIE condition in the feedback compared to the middle V1 layers appears to be superimposed upon this small, unspecific increase affecting V1 as a whole. The stronger increase in the middle compared to other layers during the FIP condition may reflect layer IV’s documented sensitivity to stimulus transients (Miller, 2003; Self et al., 2013; Zhuang et al., 2013). fMRI studies have shown that stimuli separated by time intervals of a few 100 ms evoke stronger cumulation of responses (Zhou et al., 2018) than stimuli separated by smaller time intervals. Larger time intervals between stimuli may also lead to a stronger reset of and release from inhibitory gain control (Carandini & Heeger, 2012). Layer IV with its more transient responses may be relatively more affected by the reset of gain control mechanisms and response accumulation, which may explain the stronger signal increase in the middle layers compared to feedback layers in the FIP condition.

Our finding of a neural correlate of Troxler filling-in in deep layers of V1 is in line with a large body of evidence that supports the existence of neural correlates of surface perception in early visual cortex for a number of surface features, including brightness (Komatsu et al., 2000; Malik & Boyaci, 2024; Pereverzeva & Murray, 2008; Roe et al., 2005; Salmela & Vanni, 2013; van de Ven et al., 2012), color (Hsieh & Tse, 2006, 2010; Sasaki & Watanabe, 2004) and texture (De Weerd et al., 1995; Komatsu, 2006). Some studies have shown that surface luminance variations and brightness variations have neural correlates respectively in V1 blobs and V2 thin stripes (Livingstone & Hubel, 1988; Sincich & Horton, 2005), and not in other V1 and V2 compartments related to boundary perception. This is in line with the idea that blobs and thin stripes fulfil a dedicated role in surface perception (Roe et al., 2005), and with separate modules for boundary and surface processing in computational models (Grossberg, 1987a, 1987b). Together with Kok et al. (2016), the present study is only one of two studies in humans demonstrating that neural correlates of surface perception in early retinotopic cortex involve feedback to deep cortical layers. Our data contribute to ongoing debates on the neural mechanisms of surface perception (Chan et al., 2024; Devinck & Knoblauch, 2019). In particular, our data are compatible with views that highlight the importance of recurrent interactions of retinotopic cortex with high-level areas (Gerardin et al., 2018; Perna et al., 2005; Welchman & Harris, 2003) and among retinotopic areas (Gerardin et al., 2018; Hung et al., 2007; Lee & Nguyen, 2001) during surface perception and filling-in, and inform theoretical views that link perceptual filling-in to predictive coding (Raman & Sarkar, 2016; Shipp, 2024). On the other hand, our data question broad proposals in which surface features are encoded exclusively in high-level areas in the absence of neural correlates in low-level visual areas (Dennett, 1991; von der Heydt et al., 2003; Welchman & Harris, 2003). Further work is necessary to investigate the contributions of feedback for other paradigms and surface stimuli, and to investigate how attention interacts with neural correlates of surface perception.

## Methods

### Participants

Ten healthy participants (five females) with normal or corrected to normal eyesight participated in the fMRI experiments. Participants had no history of neurological disorders. Participants provided informed consent and received study credits or monetary compensation in exchange for participation. Ethics approval was obtained from the Ethics Review Committee for Psychology and Neuroscience (ERCPN, OZL_261_148_12_2022), and all procedures followed the principles expressed in the Declaration of Helsinki.

### Stimuli

For the main fMRI experiment, three types of stimuli in the main run and separated localizer run were used (Figure 1A-C). The Filling-In Enabling (FIE) stimulus consisted of a homogeneous grey square-shaped figure (2.75° x 2.75°) placed in the middle of a dynamic texture background while participants fixated a fixation spot at the bottom right of the screen, 5.5° away from the figure. The texture was created by randomly setting non-overlapping white bars (0.06° x 0.72°) on a black background in which 21.3 % of screen was filled with the white bars. The dynamic texture field was refreshed at 30 Hz by pseudo-randomly selecting frames from 25 random versions of the texture while preventing immediate repetitions of the same pattern. The grey square and surrounding texture were equiluminant (5 cd/m2 throughout the display). The Filling-in Preventing (FIP) stimulus was identical to the FIE stimulus, except for the replacement of the texture background each second by full-field homogeneous grey for 0.3 s. The third stimulus was a full-field texture (TEX), in which only the dynamic texture was presented.

In the localizer run, we independently determined the retinotopic representation of the square figure in the visual field where we aimed to induce perceptual filling-in (Supplementary Figure S2). The stimulus consisted of a narrow square frame (thickness of 0.17°) filled with a high-contrast 16 Hz flickering checkerboard pattern. The frame expanded in size from 0.34° x 0.34° in the center of the filling-in figure to 5.84° x 5.84° well outside the figure in 17 steps at a rate of 2.53 s (1 volume)/step. We used a phase-encoding regression approach to assign voxels in and outside the figure to the different steps.

### Experimental Design

The fMRI experiments consisted of a 2-hour session comprising anatomical image acquisition, four localizer runs, and six functional runs devoted to the main experiment. Each localizer run consisted of a concatenation of 8 repetitions of the sequence of expanding frames (each sequence lasting in total 43 s), yielding a 6 min duration per run. Functional runs in the main experiment used a block design whereby 30.36 s stimulus-on periods (12 volumes) alternated with 20.24 s stimulus-off periods (8 volumes) (Figure 1D), and lasted 7 min. Each functional run presented 8 stimulus-on periods. FIE/FIP and TEX conditions were presented in independent runs. In the four runs presenting FIE and FIP conditions, four blocks per condition were shown in a counterbalanced order. In the two runs focused on the TEX condition only, eight blocks per run were presented. During each FIE and FIP block, participants were asked to rate the filling-in level numerically from 0 (perceptually no change at the figure) to 4 (completely filled-in). Level 0 was signaled by an absence of any button press, whereas levels 1-4 were signaled by pressing buttons with respectively index, middle, ring and pink fingers, see also Figure 1E) using an 8-button controller (Current designs, USA, https://www.curdes.com/hhsc-2x4-l.html). For levels 1-4, participants were instructed to start pressing the corresponding key once a specific level of filling-in was perceived, and to keep that key pressed until the level of filling-in changed. Participants were furthermore instructed to then release the key and press another key (or stop pressing any key) in line with their next filling-in percept. All participants who participated in the fMRI experiment had participated previously in behavioral pilot experiments to verify that they showed filling-in in the FIE condition, and a lack of filling-in in the FIP condition. We required that the average filling-in level throughout the trial and across trials in the FIE condition would be at least twice the one in the FIP condition for participants to be included in the experiment. From the 33 participants we tested in pilot psychophysical experiments, we scanned best 10 fillers who passed this criterion and agreed to participate in the fMRI experiment. The behavioral data distribution across participants in the pilot experiment is presented in Supplementary Figure S5. Prior to these pilot and fMRI experiments, participants were exposed to the images on the screen illustrated in Figure 1E to help them associate the numeric responses to specific stages of filling-in. Following the fMRI experiments, participants responded positively to the question whether the illustrations of filling-in levels corresponded well to the stages they experienced during perceptual filling-in.

### Scanning

We used a Siemens MAGNETOM 7T Plus scanner with a 1TX/32RX head coil at Scannexus (Maastricht, NL). For functional and localizer runs, we collected 2D-GE-BOLD data with a resolution of 0.8 mm isotropic, 66 slices, a TR of 2.53 s, a TE of 26.4 ms, flip angle of 67°, a multi-band acceleration factor of 2, a GRAPPA acceleration factor of 3, and a partial Fourier of 6/8. Although we acquired functional images with an anterior-posterior (AP) phase encoding direction, five volumes of PA images were acquired in each session for distortion correction. For anatomical imaging, an MP2RAGE protocol with a resolution of 0.7 mm isotropic and TA of 8 min was used. B0-shimming was optimized using procedures described in (Tse et al., 2016).

### Analysis

#### Preprocessing

BrainVoyager (version 22.4) (Brain Innovation, Maastricht, NL) was used for fMRI data analysis (Goebel et al., 2006). Standard preprocessing (slice scan time correction, 3D motion correction, and high-pass filter with GLM-Fourier 6 cycles) was applied. Distortion correction was done with FSL (version 6.0) and a top-up obtained from www.nitrc.org (Jenkinson et al., 2012) using AP and PA images that were acquired in each session. Following distortion correction, anatomical and functional images were registered on BrainVoyager using boundary-based registration (BBR), and further analysis was done in the surface space.

### Cortical segmentation and layerification

We used the “Advanced segmentation” tool in BrainVoyager to perform cortical segmentation. The default parameters were applied for each step of the tool. To enhance the quality of the segmentation, intermediate anatomical data were manually corrected after masking subcortical structures and removing the cerebellum. Then, manual segmentation was done on the final grey matter (GM) and white matter (WM) segmentation, specifically in our region of interest. All manual segmentation was done on ITK-SNAP (version 3.8.0) obtained from www.itksnap.org (Yushkevich et al., 2006) with a Wacom drawing pad (https://www.wacom.com, Kazo, Saitama, Japan) after converting to Nifti files by bvbabel (https://github.com/ofgulban/bvbabel). From 11 mesh time-course files (surface data) across depth within GM, we included meshes 1-3 into deep, 5-7 into middle, and 9-11 into superficial layer compartments (see Supplementary Figure S5). For the whole-GM analyses within a region-of-interest (ROI) definition, we included all 11 mesh time-course files.

### Localizer Analysis

We identified the cortical representation of the square figure (region of interest, ROI) by averaging all four runs of the localizer data in each session and then calculated linear correlation using the “compute linear correlation map” command in BrainVoyager to obtain a retinotopic map of the ROI on the surface. We did not perform whole-field retinotopic mapping but determined the location of an ROI as belonging to V1 or V2 based on anatomical criteria. Given the presentation of the square figure was always in the upper left quadrant of the visual field relatively close to the horizontal meridian, the V1 ROI appeared in the lower bank of the calcarine sulcus, and the extrastriate on the lower gyrus of the calcarine sulcus. We successfully determined a V1 ROI in all participants but determined an extrastriate ROI in only six of ten participants (see Supplementary Figure S2). However, in those 6 participants we found that the extrastriate ROIs were too small for further analysis and for separating the cortex within the ROI into layers. We therefore focused our analysis on striate cortex.

### Functional Analysis

After all preprocessing steps, we computed event-related averages for each condition on the cortical surface. As a baseline for the event-related averages, we used three data points preceding the start of each block, and averaged all three values to obtain a baseline value for each block. After subtracting baselines from each block, we calculated the mean across all blocks for each condition for each participant, and reported the mean and standard error across participants (Supplementary Figure S3).

For subtraction analyses, we first converted the time course of each trail into z-scores using the 3 data point prior to the stimulus onset to 13 data point after stimulus onset. Early and late windows were defined by taking the mean value of 5 time points from the 3rd data point, reflecting from 7.59 s to 17.71 s after stimulus onset (early window), and from the 9th data point, reflecting from 22.77 s to 32.89 s (late window). The subtraction analysis was done by subtracting the averaged value in the early period from the late period. For the supplementary time point analysis, both windows were defined by fetching 4 time points, starting from taking the 3^rd^ data point and shifting time points for early windows, while fixing the late window definition (25.3 s - 32.89 s).

For layer analysis, we defined our ROI by calculating the overlaps between localized ROI and the anatomically defined cortical layer groups (see the cortical segmentation and layerification above for the details). Subtraction was done in the same manner as in the whole GM analysis (see above).

## Acknowledgement

This work was supported by Nederlandse Organisatie voor Wetenschappelijk Onderzoek (NWO) Open Competition grant 406.21.GO.044 to PDW. We acknowledge Desmond H.Y. Tse and Scannexus for scanning and advance shimming, Mario Senden for a discussion of the localizer paradigm, and Omer Faruk Gülban for consultation regarding layer analysis.

## Notes

### Competing Interest Statement

The authors have declared no competing interest.

