## Supplementary Figures for "Feedback to deep layers in human V1 during perceptual filling-in"

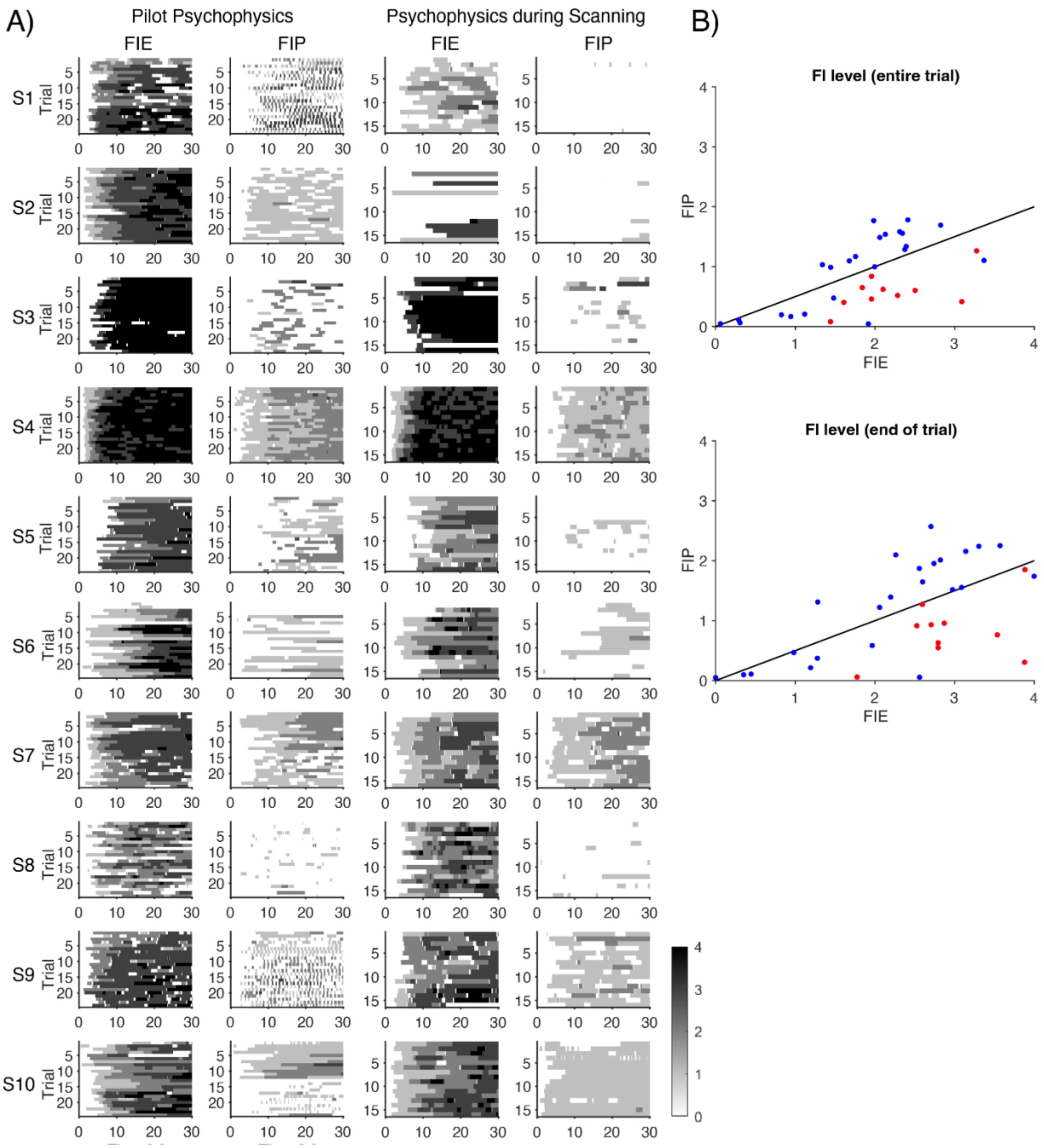


*Supplementary Figure S1. Filling-in reports from pilot psychophysical experiment and fMRI experiment. (**A**) Individual participant data showing pilot psychophysics collected outside the scanner (lefthand two columns) and psychophysics from the same participants collected during scanning (righthand two columns). Each graph shows the level of filling-in (darker grey levels symbolize stronger filling-in) as a function of time per trial number. Note the stronger filling-in in the filling-in enabling (FIE) compared to the filling-in prevention (FIP) condition, and the consistency of results between the behavioral data collected outside and inside the scanner. The behavioral data from participant S2 collected in the scanner were not included in the group average (Figure 2B, C) because a post-experiment check showed that several buttons used to signal the filling-in level hadn't functioned reliably. Because participant S2 had shown*

*reliable filling-in in the FIE condition and limited filling-in in the FIP conditions during pilot psychophysics, and because the limited data collected in the scanner coincided with that pattern, we kept participant S2 included in the fMRI sample. Statistical analysis outcomes of tests applied to psychophysical and fMRI data (in terms of reaching significance) did not depend on whether participant S2 was included or not. (**B**) Psychophysical pilot filling-in data in FIE and FIP conditions in a sample of 33 participants. Filling-in levels in the FIP condition were plotted as a function of filling-in levels in the FIE conditions after averaging data across the total trial duration (top panel) or after averaging data from the last 5 s of the trial only (bottom panel). Participants for the fMRI experiment were selected based on availability from the subgroup showing at least twice as much filling-in in FIE compared to FIP conditions based on whole-trial and late-trial analyses (10 red dots falling below the black criterion line).*

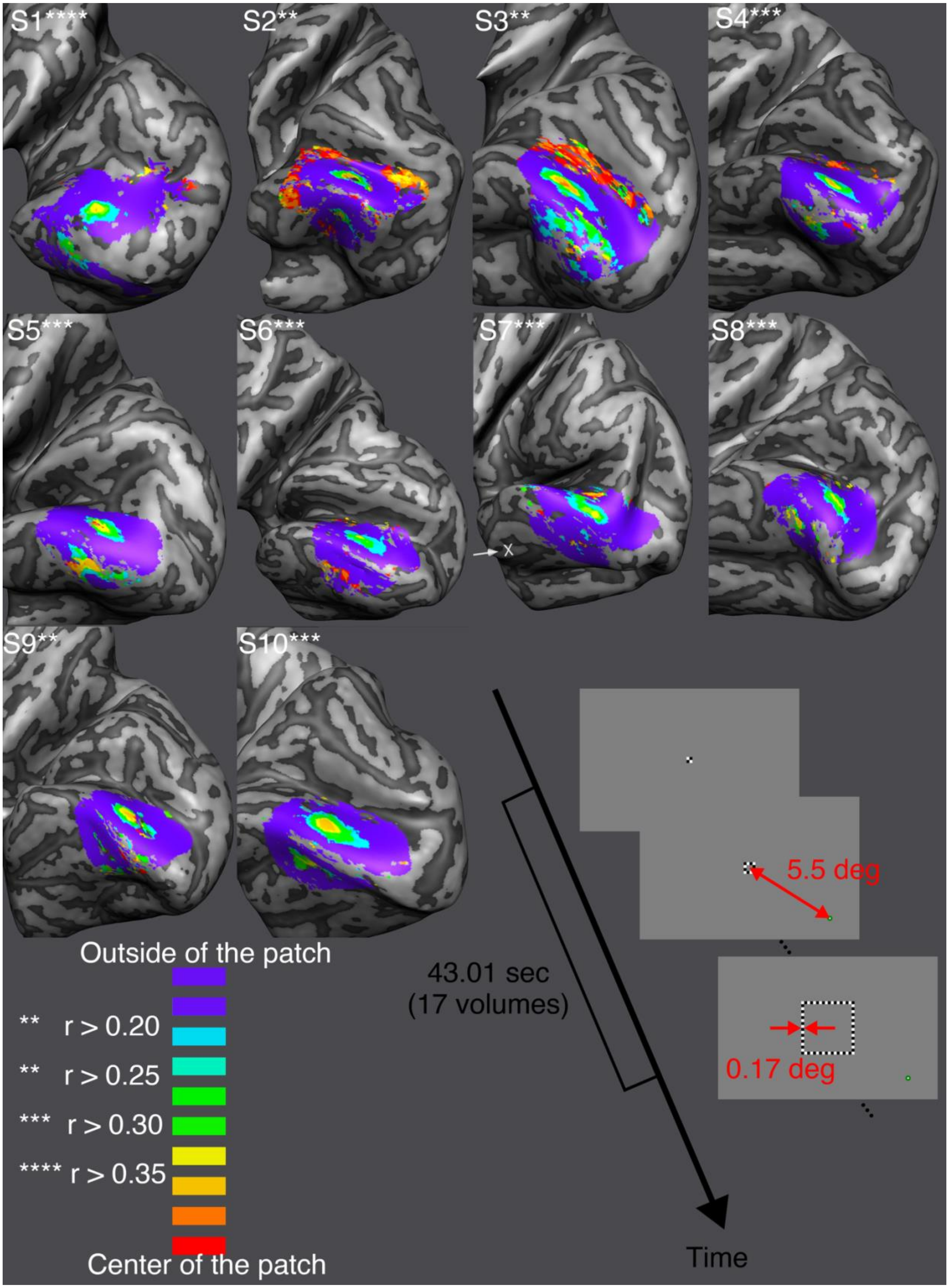

S1****
S2**
S3**
S4***
S5***
S6***
S7***
S8***
S9**
S10***
X
Outside of the patch
** r > 0.20
** r > 0.25
*** r > 0.30
**** r > 0.35
Center of the patch
43.01 sec
(17 volumes)
5.5 deg
0.17 deg
Time

*Supplementary Figure S2. Individual localizer surface maps and phase encoding localizer design. Color shows phase-encoded steps of the localizer frame (see Methods) regressed by a linear correlation model. Any color other than purple represents the inside of the figure whereas purple represents the outside of the figure. Thin square localizer frames filled with a 16 Hz flickering checkerboard texture were centered on the middle of the figure and increased in size from small (0.34° x 0.34°) to large (5.84° x 5.84°) in 17 steps (8 steps within the figure) that each took a duration of one volume (2.53 s). The frame dimensions represent the outer border of the frames, which had a thickness of 0.17°.*

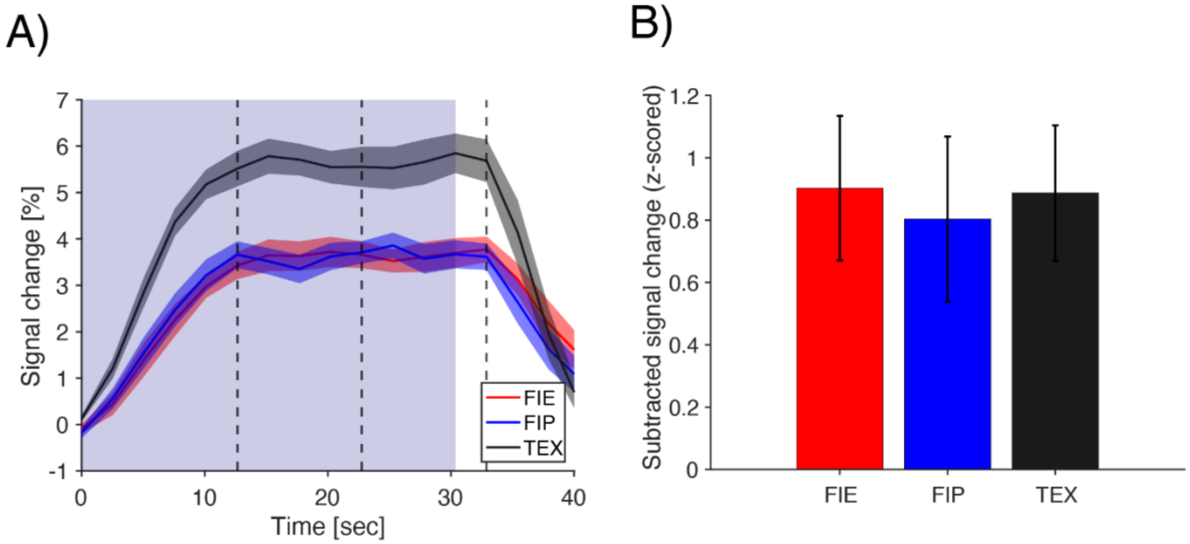


*Supplementary Figure S3. fMRI data in the filling-in enabled (FIE), filling-in prevention (FIP), and full texture (TEX) conditions.* ***(A)*** *Averaged (across participants) fMRI signal elicited in the three conditions of interest. Shading shows standard error of the mean across participants. The blue background represents stimulus duration.* ***(B)*** *Average (across participants) fMRI response in later compared to earlier time windows for the three conditions of interest (FIE, FIP, TEX). Positive values indicate a larger average response the later time window. Error bars represent the standard error of the mean. An ANOVA revealed a main effect of time, without a time x condition interaction (time, $F(1,36) = 21.47$, $p < 0.001$; condition, $F(1,36) = 0.54$, $p = 0.46$; interaction, $F(1,36) = 0.12$, $p = 0.72$). Furthermore, laminar analysis showed a positive increase for all layers in FIE (paired t-test $t(9) = 5.92$, $p = 0.001$ in deep, $t(9) = 4.26$, $p = 0.012$ in middle, $t(9) = 4.04$, $p = 0.018$ in superficial layers), and in FIP ($t(9) = 3.03$, $p = 0.085$ in deep, $t(9) = 3.69$, $p = 0.030$ in middle, $t(9) = 3.55$, $p = 0.037$ in superficial layers). All p-values underwent Bonferroni correction.*

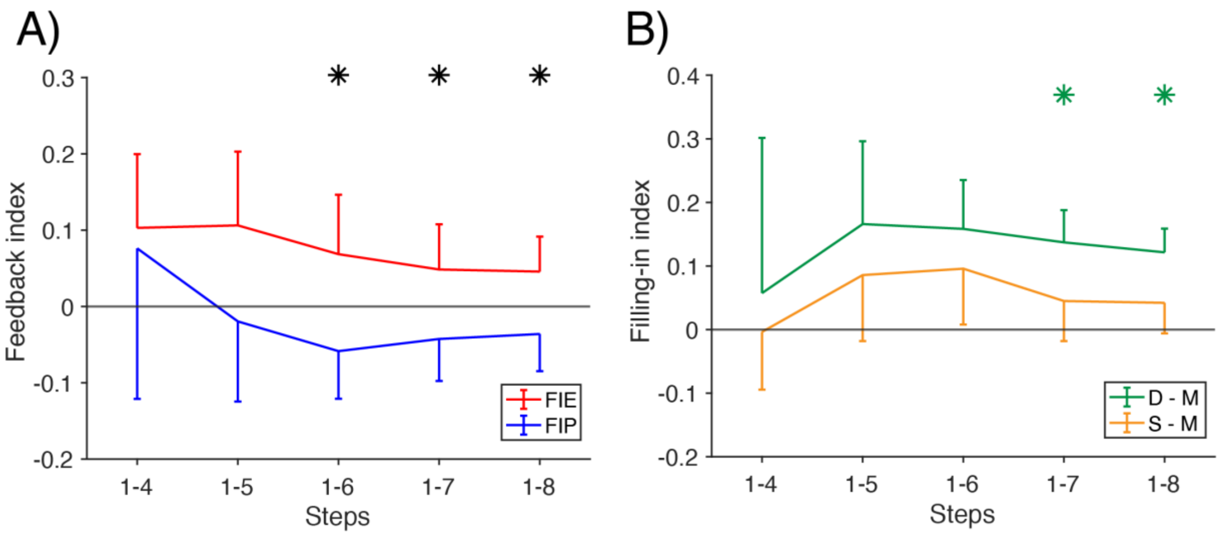


*Supplementary Figure S4. ROI selection analysis.* ***(A)*** *Feedback index for different selections of vertices within the ROI of the figure. Solid lines indicate means and error bars indicate standard errors of the mean across participants. Black stars indicate statistically significant differences in feedback index between filling-in enabling (FIE) and filling-in prevention (FIP) conditions (paired t-test, $p < 0.05$).* ***(B)*** *Filling-in index contrasted between deep and middle layers (D-M) or between superficial and middle layers (S-M) for different selections of vertices within the figure ROI. Green stars indicate D-M differences that deviated statistically from zero (paired t-test, $p < 0.05$). There were no significant S-M differences. Other conventions as in A.*

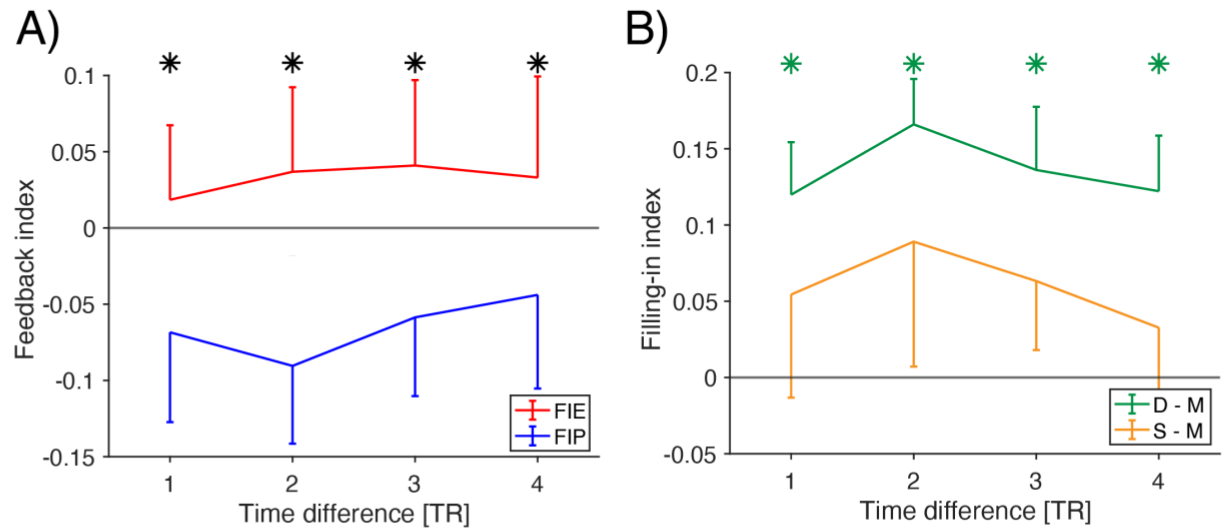


*Supplementary Figure S5. Time window analysis.* ***(A)*** *Feedback index for time differences between early and late window definitions ranging from 1 to 4 TRs (one TR = 2.53s). The time difference on the x-axis represents the difference between the latest time point of the early window and the earliest time point of late window. Solid lines indicate means and error bars indicate standard errors of the mean across participants.*

*Black stars indicate statistically significant differences in feedback index between filling-in enabling (FIE) and filling-in prevention (FIP) conditions (paired t-test, $p < 0.05$).* ***(B)*** *Filling-in index contrasted between deep and middle layers (D-M) or between superficial and middle layers (S-M), and normalized, for different time intervals between early and late windows used for computing signal changes within the figure ROI. Green stars indicate D-M differences that deviated statistically from zero (paired t-test, $p < 0.05$). There were no significant S-M differences. Other conventions as in A.*
